## Supplementary figures for "Comparison of sequence- and structure-based antibody clustering approaches on simulated repertoire sequencing data"

### Supporting figures

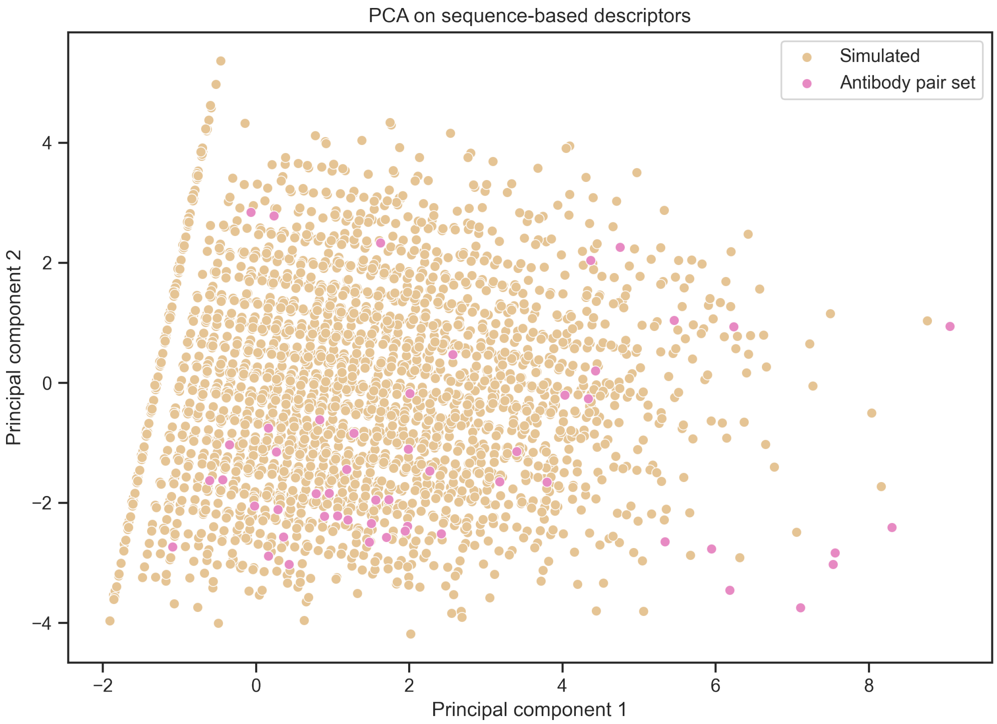


**Fig 1. PCA of full simulated repertoire using sequence descriptors.** A PCA was fitted using sequence descriptors of the full simulated repertoire and the antibody pair set. The first and second principal component are shown. The annotated antibodies (pink) overlap strongly with the simulated clones (beige), indicating that these antibody sets are not consistently different from each other.

A

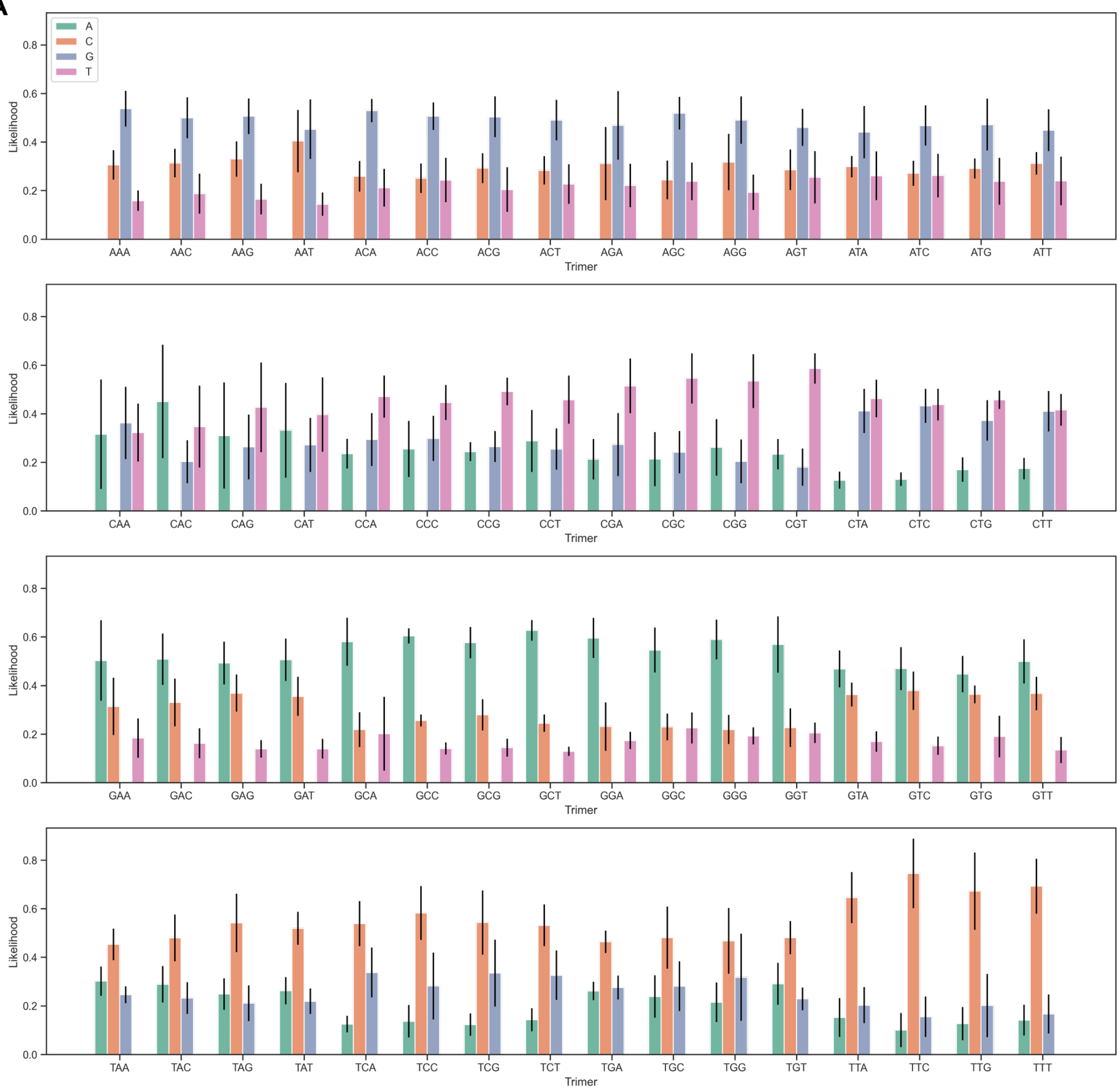

A
C
G
T
Likelihood
Trimer
AAA
AAC
AAG
AAT
ACA
ACC
ACG
ACT
AGA
AGC
AGG
AGT
ATA
ATC
ATG
ATT
CAA
CAC
CAG
CAT
CCA
CCC
CCG
CCT
CGA
CGC
CGG
CGT
CTA
CTC
CTG
CTT
GAA
GAC
GAG
GAT
GCA
GCC
GCG
GCT
GGA
GGC
GGG
GGT
GTA
GTC
GTG
GTT
TAA
TAC
TAG
TAT
TCA
TCC
TCG
TCT
TGA
TGC
TGG
TGT
TTA
TTC
TTG
TTT

B

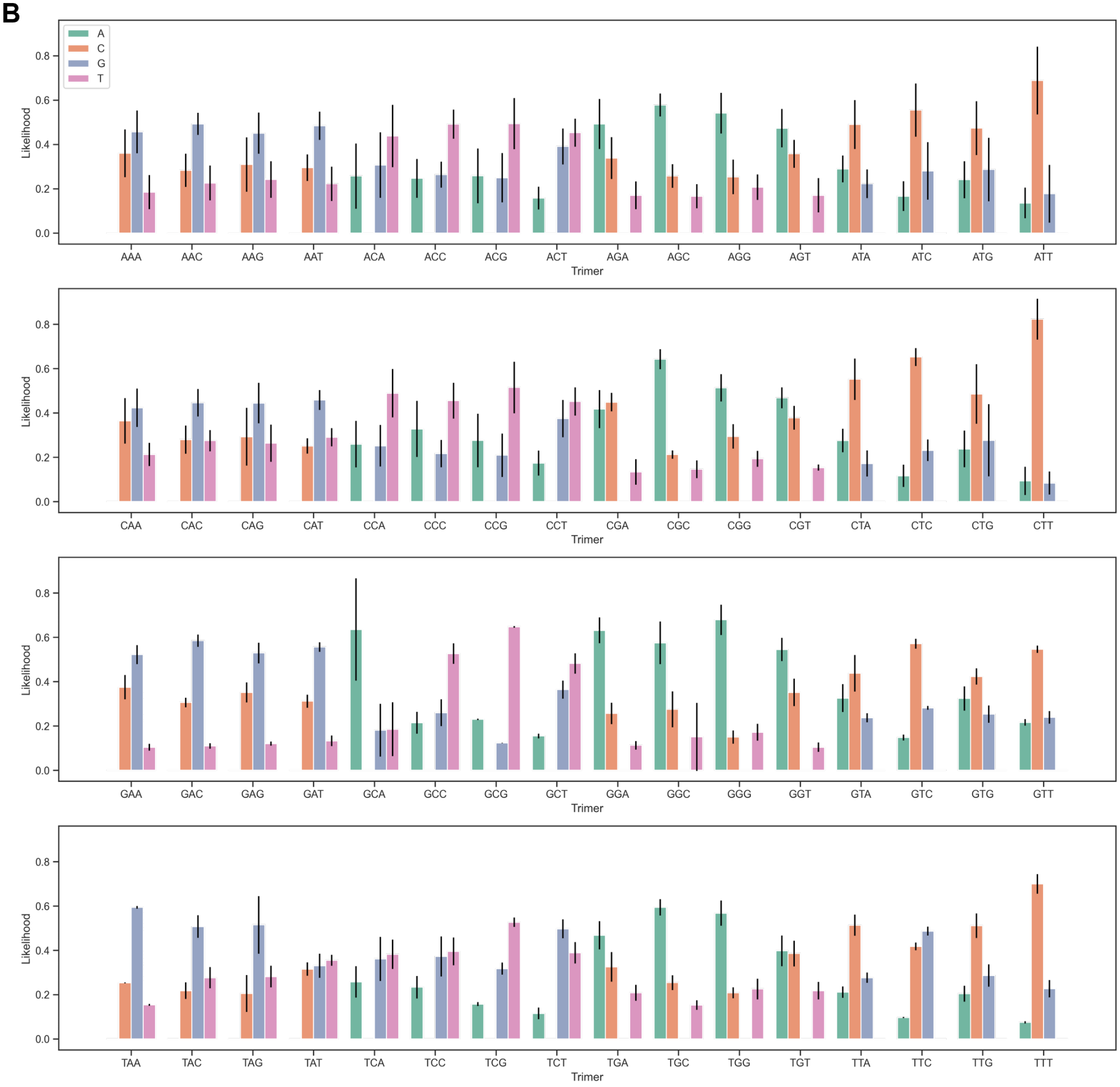
A
C
G
T
Likelihood
Trimer
AAA AAC AAG AAT ACA ACC ACG ACT AGA AGC AGG AGT ATA ATC ATG ATT
CAA CAC CAG CAT CCA CCC CCG CCT CGA CGC CGG CGT CTA CTC CTG CTT
GAA GAC GAG GAT GCA GCC GCG GCT GGA GGC GGG GGT GTA GTC GTG GTT
TAA TAC TAG TAT TCA TCC TCG TCT TGA TGC TGG TGT TTA TTC TTG TTT

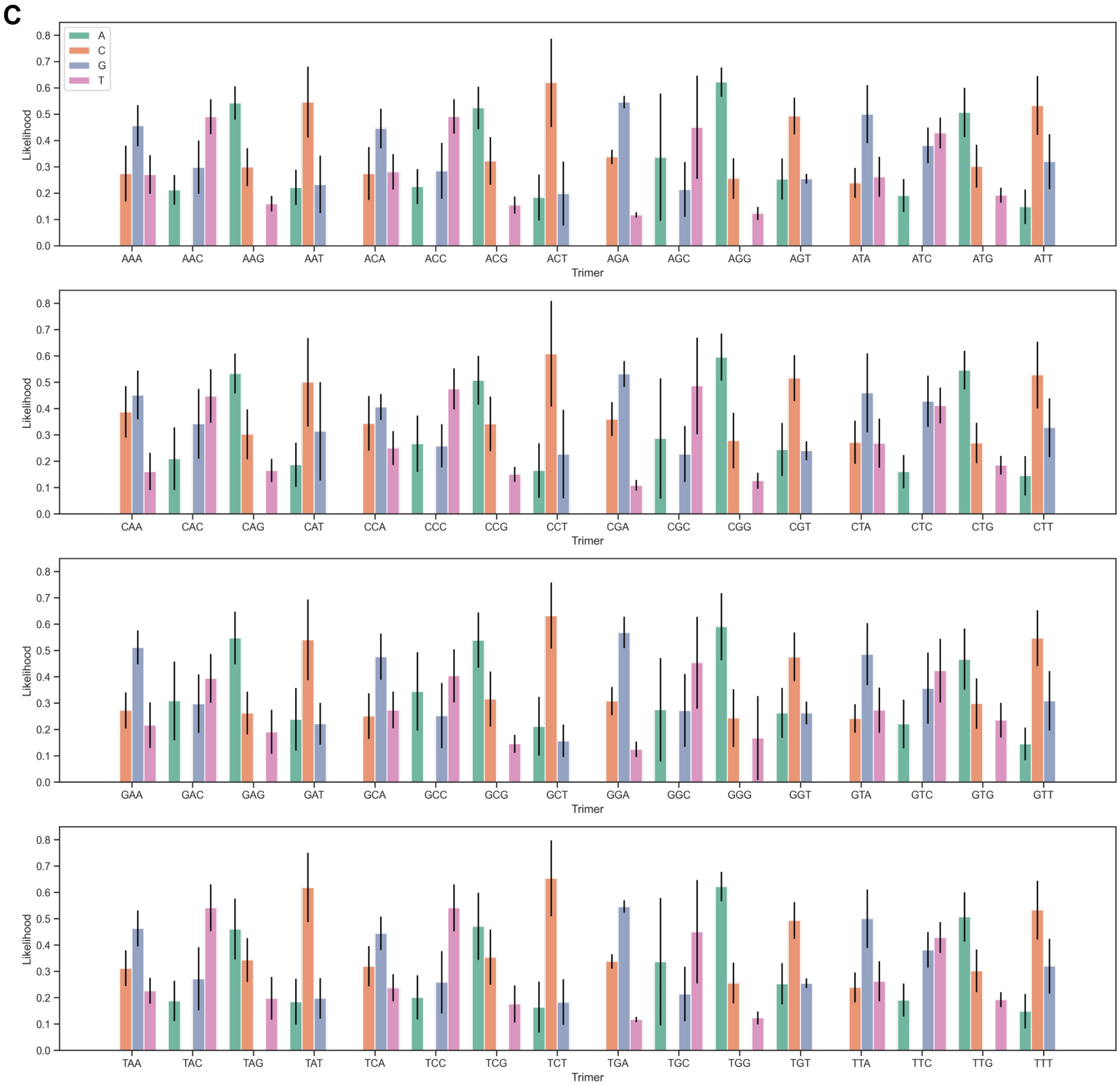


**Fig 2. Aggregated substitution likelihoods.** Observed and inferred substitution likelihoods of single nucleotides within immune receptor sequences have been used for backtranslation of amino acid to nucleotide sequences of the curated antibodies. The substitution likelihoods were provided for fivemers. To simplify backtranslation these fivemer substitution likelihoods were aggregated to trimer substitution likelihoods. The aggregated likelihoods for each substitution including standard deviations are shown for the start (A), center (B) and end (C) nucleotide of each possible trimer.
